# The urban-adapted underground mosquito, *Culex molestus*, maintains exogenously influenced circadian rhythms despite an absence of photoperiodically induced dormancy

**DOI:** 10.1101/2020.10.02.323824

**Authors:** Natalie R. Epstein, Kevin Saez, Asya Polat, Steven R. Davis, Matthew L. Aardema

## Abstract

In temperate climates, the mosquito *Culex molestus* lives almost exclusively in urban underground locations such as flooded basements, sewer systems and subway tunnels. Unlike most other mosquito taxa found at higher latitudes, *Cx. molestus* remains active and continues to breed throughout the winter. This is attributable to year-round above freezing temperatures in its preferred underground habitats combined with an inability to enter a state of arrested development (‘diapause’) in response to shortening photoperiods in autumn. Prior studies have shown that the genes associated with circadian rhythms (i.e. ‘clock genes’) also influence the photoperiodic induction of diapause in the closely related mosquito, *Cx. pipiens*. These results suggest that molecular changes in one or more clock genes could contribute to the absence of photoperiodically induced diapause in *Cx. molestus*. As *Cx. molestus* predominantly inhabits underground locations generally devoid of a predictable photoperiod, such mutations may not be removed by purifying selection as they would have minimal fitness consequences. To examine the possibility that *Cx. molestus-* specific genetic changes in one or more clock genes correlate with its inability to enter a photoperiodically induced dormant state, we first used genomic data to search for inactivating mutations or other structural variants in genes known to influence circadian rhythms in Diptera (‘flies’). We further investigated non-synonymous, derived genetic divergence in this same class of genes. Next, we generated transcriptome data from multiple life-stages of *Cx. molestus* to survey binary expression of annotated clock genes throughout this mosquito’s lifecycle. Finally, we carried out experimental studies to assess the extent to which *Cx. molestus* retains exogenously influenced circadian rhythms, and whether it harbors any tendencies towards dormancy when exposed to a shortened photoperiod. Our results indicate that the gene Helicase domino (*dom*) has a nine-nucleotide, in-frame deletion specific to *Cx. molestus*. Previous work has shown that splice variants in this gene influence circadian behavior in *Drosophila melanogaster*. We also find derived, non-synonymous single nucleotide polymorphisms (SNPs) in eight genes that may also affect circadian rhythms and/or diapause induction in *Cx. molestus*. Four other circadian genes were found to have no quantifiable expression during any examined life stage, suggesting potential regulatory variation. Our experimental results confirm that *Cx. molestus* retains exogenously-influenced circadian rhythms but is not induced to enter a dormant state by a shortened photoperiod. Collectively, these findings show that the distinct, but potentially molecularly interconnected life-history traits of diapause induction and circadian rhythms are decoupled in *Cx. molestus* and suggest that this taxon may be a valuable tool for exploring exogenously influenced phenotypes in mosquitoes more broadly.

## INTRODUCTION

Examining the ways in which distinct life-history traits are linked to one another can provide a better understanding of the mechanisms through which organisms adapt to, or are constrained by, environmental conditions (Blows and Hoffmann, 2005; Mauro and Ghalambor, 2020). In insects, two important life-history traits that are tightly linked to environmental variation are circadian rhythms and seasonal dormancy in response to prolonged, adverse climatic conditions (Tauber and Tauber, 1981; Danks, 1987; Kyriacou et al., 2008). Individual variation in both of these traits can have substantial fitness consequences, with mis-timed behavioral responses leading to reduced or absent mating opportunities, reduced access to food resources, or increased probability of mortality (Danks, 1987; Yerushalmi and Green, 2009).

For both circadian rhythms and seasonal dormancy, the perception of external light cues plays an important role in their initiation and/or maintenance (Beck, 1980; Denlinger et al., 2017). Insect circadian rhythms are usually exogenously entrainable, with the daily succession of light and dark influencing many biological processes that oscillate over a 24-hour period. This entrainability allows individuals to adjust their behavior to match seasonal or geographic variation in day-length (Meireles-Filho and Kyriacou, 2013). A dormancy response to predictable seasonal changes, such as colder periods in temperate climates (i.e. ‘winter’), is also often strongly influenced by changes in daylength (Adkisson, 1966). In insects, one of the most common responses to oncoming inclement weather that is seasonally consistent is diapause. Insect diapause is a state of dormancy in which growth and development is halted until the environment changes to more favorable conditions (Tauber and Tauber, 1976). It is common in insects living in temperate regions where the summers are warm with long days and the winters are cold with short days (Denlinger, 2002).

Mosquitoes of the *Culex pipiens* species complex are important disease vectors of many arboviruses including West Nile virus and Saint Louis encephalitis (Fonseca et al., 2004). For this reason, understanding the nature and manifestation of circadian rhythms and seasonal dormancy in these taxa has implications for vector population control and disease mitigation (Ewing et al., 2019). In one member of this complex, the Northern House Mosquito (*Culex pipiens*), only females enter diapause, spending the winter remaining relatively stationary in sheltered locations such as tree holes or natural overhangs (Vinogradova, 2000). These diapausing females experience no growth in their ovarian follicles and will not seek out a blood meal until spring and the termination of diapause (Denlinger and Armbruster, 2014). Like most diapausing insects, they rely on their metabolic reserves stored prior to entering the diapause state (Zhou and Miesfeld, 2009). Male *Cx. pipiens* do not diapause and die with the onset of winter. *Cx. pipiens* populations go through multiple generations each year, and as such, diapause induction is facultative, being initiated by the shorter photoperiods and cooler temperatures experienced by larvae in late summer and autumn (Eldridge, 1968; Spielman and Wong, 1973).

As both circadian rhythms and the initiation of diapause are closely tied to external light cues, it is perhaps not surprising that they are correspondingly linked molecularly in *Cx. pipiens*, with many of the same genes influencing both life-history traits (e.g. Ikeno et al., 2010; Meuti and Denlinger, 2013). In females, functioning clock genes are required to initiate diapause (Meuti et al., 2015). Under long day, diapause-averting conditions, the suppressed expression of the genes period (*per*), timeless (*tim*), and cryptochrome2 (*cry2*) results in an elevated expression of pigment dispersing factor (*PDF*), which has a role in circadian timing and diapause induction. However, if *PDF* expression is suppressed in females reared in long day conditions, the diapause phenotype occurs. In short day, diapause-inducing conditions, the expression of *per, tim*, and *cry2* is elevated and the diapause phenotype occurs (Hand et al., 2016).

*Cx. pipiens*’ broadly co-occurring sister taxon, *Cx. molestus*, is an urban adapted mosquito found predominately in cities where it lives in subterranean locations such as flooded basements, sewers and subway tunnels (Byrne and Nichols, 1999; Vinogradova, 2000). Both observational and experimental studies indicate that, unlike *Cx. pipiens*, this mosquito lacks an ability to enter a diapause state in response to a shortened photoperiod (Richards, 1941; Spielman and Wong, 1973; Kassim et al., 2013; Nelms et al., 2013; Bajwa and Zuzworsky, 2016). The taxonomic status of *Cx. molestus* remains a challenging biological problem, and many authors treat it either as a subspecies or ecological form of *Cx. pipiens* (e.g. Vinogradova, 2000; Harbach, 2012). However, in addition to their important difference in diapause ability, they also differ in host preferences, mating behaviors, and the requirement of a blood meal prior to egg-laying (Fonseca et al., 2004). With regards to this last characteristic, one of the predominant distinguishing characteristics of *Cx. molestus* is an obligate need to lay one batch of eggs prior to seeking a vertebrate host (‘autogeny’; Spielman, 1971). For the sake of simplicity, in this study we will refer to each taxon as *Cx. molestus* and *Cx. pipiens* respectively, with no implied taxonomic status intended.

In one study examining circadian rhythms in *Cx*. molestus, the authors concluded this taxon displays less sensitivity to photoperiod than other members of the genus (Chiba et al., 1981). In another study, it was shown that adult eclosion (the emergence of the adult from the pupal state) of *Cx. molestus* populations from Russia are less influenced by exogenous factors than are *Cx. pipiens* populations (Karpova, 2009). Finally, a third study showed that adult *Cx. pipiens* females from the United States displayed more flexibility in when they would take a blood meal compared to *Cx. molestus* populations (Fritz et al., 2014). The authors of this study suggested this may be explained by differences in response to photoperiod displayed by these two mosquito taxa. Taken together, these studies strongly support the hypothesis that exogenous entrainment of circadian rhythms via light cues may be diminished in *Cx. molestus*. As *Cx. molestus* inhabits predominantly underground locations, populations are unlikely to experience natural fluctuations in photoperiod. It is therefore possible that potential selection pressures to maintain a response to external photoperiod cues has been reduced or eliminated. This, in turn, could also have led to a potential loss in diapause ability.

In this study, we investigate the presence of *Cx. molestus*-specific genetic changes that may be associated with its inability to express a photoperiodically induced dormant state. Specifically, we postulate that an inactivating mutation or other major structural variant in one or more clock genes could simultaneously account for its inability to enter diapause and a likely reduction or loss of entrainable circadian rhythms. We also look for non-synonymous single nucleotide polymorphism (SNPs, aka ‘missense mutations’) in these genes, and use expression data to survey possible regulatory influences. Finally, we experimentally examine photoperiodically entrainable circadian rhythms and diapause induction in a *Cx. molestus* population originating from New York City, postulating that both life-history traits are reduced or absent in this mosquito.

## MATERIALS AND METHODS

### Identifying genes that may contribute to circadian rhythms and/or diapause

One of the primary goals for this study was to identify *Cx. molestus*-specific genetic variation in genes potentially influencing circadian rhythms and diapause. To facilitate this, we took advantage of the assembled and annotated genome of the closely related mosquito, *Cx. quinquefasciatus* (VectorBase assembly CpipJ2; Arensburger et al., 2010). This taxon, the ‘Southern House Mosquito’, is a member of the *Culex pipiens* species complex (as are *Cx. molestus* and *Cx. pipiens*), and its annotated genome is frequently used in genetic and genomic comparisons between *Cx. molestus* and *Cx. pipiens* (e.g. Asgharian et al., 2015; Price and Fonseca, 2015; Aardema et al., 2020; Yurchenko et al., 2020). We first needed to determine which annotated *Cx. quinquefasciatus* genes potentially influence circadian rhythms and/or diapause. To do this, we used the search words ‘circadian’, ‘photoperiod’, ‘dormancy’, ‘light stimulus’ and ‘diapause’ to locate GO categories associated with these terms in *Drosophila melanogaster* (Flybase, Dmel Release 6.32; Thurmond et al., 2019). The specific GO terms found are listed in Table S1. Next, we determined all *D. melanogaster* genes that had a ‘biological process’ associated with one or more of these terms. The *D. melanogaster* peptide sequences for these genes were compiled into a dedicated FASTA file. Using a local protein-protein BLAST v. 2.7.1 (‘blastp’; Altschul et al., 1990), and these *D. melanogaster* sequences as our queries, we searched the annotated peptide sequences of *Cx. quinquefasciatus*, with an evalue of 1e-10 and default values for all other settings. We considered the ‘best’ matches (the longest sequence with the highest percent similarity) between *D. melanogaster* and *Cx. quinquefasciatus* as prospective orthologs, provided they had greater than 50% amino acid similarity across 100 or more amino acids. This list of potential circadian genes in *Culex* contained 154 unique annotated sequences (Table S2), and it is these genes that we utilized in subsequent analyses (see below). Henceforth, these genes will be referred to as the ‘*Culex* circadian genes’.

### Structural variants in genes influencing circadian rhythms

We wanted to determine if one or more *Culex* circadian genes harbored *Cx. molestus-*specific major structural variants or other potentially ‘inactivating’ mutations that could correlate with its insensitivity to photoperiod in the induction of a dormancy state. To do this, we mapped previously published *Cx. molestus* genomic short-read Illumina data (NCBI-SRA accession numbers: SRR10053379, SRR10053380, and SRR10053386) to the *Cx. quinquefasciatus* genome (Arensburger et al., 2010) using the ‘MEM’ algorithm implemented in the program BWA v. 0.7.15 (Li, 2013) with default settings. The sample used to produce these genomic reads came from a New York City-derived lab strain of *Cx. molestus* (Aardema et al., 2020). After mapping, we marked duplicate reads with the MarkDuplicates function in Picard v. 1.77 (http://broadinstitute.github.io/picard/), then performed indel realignment using the IndelRealigner function of the Genome Analysis Toolkit v. 3.8 (‘GATK’; McKenna et al., 2010). Next, we determined genotype likelihoods using the ‘mpileup’ command in bcftools v. 1.9 (Li et al., 2009), then identified sites divergent from the reference using the ‘call’ command, also with bcftools. These divergent sites included both single nucleotide polymorphism (SNPs) and insertion/deletions (INDELs). We then used the program SnpEff v. 4.3 (Cingolani et al., 2012) with a custom database for the gene annotations of *Cx. quinquefasciatus* to annotate these variants with default parameters. We cross-referenced the gene summary produced by the SnpEff program with our list of 154 *Culex* circadian genes, identifying those on the list that were determined to have a ‘high’ impact variant. Potential high impact variants included frameshift mutations, loss of the start codon, gain of a stop codon, loss of the stop codon, or disruption of an exonic splice site. This cross-referencing generated 33 *Culex* circadian genes that had one or more potential structural variants or other mutations of great effect.

To test whether these variants were valid (i.e. not an artifact of incorrect annotation), we wanted to determine that the expressed coding gene structure in *Cx. molestus* matched that of the *Cx. quinquefasciatus* gene annotation. We also wanted to assess if the variant(s) were both found in a broad geographic representation of *Cx. molestus* samples, and were not present in the sister taxon, *Cx. pipiens*. To do this, we *de novo* assembled two *Cx. molestus* transcriptomes (from samples deriving from the United States and Germany, respectively [Table S3]) and two *Cx. pipiens* transcriptomes (also from samples deriving from the United States and Germany), using the program Trinity v. 2.8.4 with default parameters (Haas et al., 2013). We performed a nucleotide-nucleotide BLAST v. 2.7.1 (‘blastn’) with the target exon containing the potential inactivating mutations as our query sequence and each of these four assembled transcriptomes as our database. Using RNA-seq (transcriptome) data allowed us to simultaneously confirm accurate annotation of the gene and that the focal variant was present in both North American and European *Cx. molestus*, but not in the closely-related *Cx. pipiens*.

If our analysis from these four transcriptomes supported a correct annotation of the focal gene and additionally if the potential ‘large-effect’ variant was present in both *Cx. molestus* samples and neither *Cx. pipiens* sample, we then examined its presence/absence in four additional *de novo* assembled genomes from two *Cx. molestus* samples (one from Belarus and the aforementioned NYC sample; see Table S3), and two *Cx. pipiens* samples (one from Belarus and one from New Jersey; see Table S3). These *de novo* assemblies were done with the program ABySS v. 2.2.3 (Jackman et al., 2017), with K values from 56 to 96 at 10 nucleotide intervals. The ‘best’ assembled genome was chosen as the K value with the highest E-size (a measure of probable gene completeness; Lian et al., 2014), as determined with ‘abyss-fac’. Again, we used a nucleotide-nucleotide BLAST v. 2.7.1 (‘blastn’) with the target exon containing the potential inactivating mutations as our query sequence and each of these four assembled genomes as our database.

### Genetic Divergence Between Diapausing and Non-Diapausing *Culex* Forms

In addition to possible structural variants in genes known to influence diapause and circadian rhythms, it is possible that derived amino acid changes could also impact the expression of these traits. To examine this possibility, we took advantage of previous research comparing non-synonymous divergence (Ka) between New York City *Cx. molestus* and a *Cx. pipiens* population from Germany (Price and Fonseca, 2015). We first compared our *Culex* circadian gene list to those which were found to have a Ka value greater than zero (indicating non-synonymous divergence between *Cx. molestus* and *Cx. pipiens*). From this comparison, we derived a list of 49 gene candidates to investigate potential derived amino acid changes in *Cx. molestus*.

Next, we used a local nucleotide-nucleotide BLAST v. 2.7.1 (‘blastn’) to compare each exon within each of these 49 genes from the annotated *Cx. quinquefasciatus* genome to each of the four previously described *de novo* genome assemblies (two *Cx. molestus* samples and two *Cx. pipiens* samples; see above). We aligned these four focal sequences with the *Cx. quinquefasciatus* focal exon and the full gene sequence to maintain the correct reading frame (in relation to the annotation). Sequence gaps, codons that spanned an intron and regions that were not present in all four focal genomes were removed. After all exon regions were located, aligned and trimmed, these sequences were concatenated into a single sequence, then converted to amino acids. Alignment, trimming and amino acid conversion was done with SeaView v. 4.6.3 (Gouy et al., 2010). Using a custom Perl script, we counted the number of derived amino acid changes observed in each genome (relative to *Cx. quinquefasciatus*).

### Expression of Clock Genes

To assess the presence of potential inactivating mutations in *Culex* circadian genes, we generated RNA-seq data for four Cx. *molestus* life stages: larva, pre-pupa, pupa and adult. For each library we combined 10 individual heads, then extracted mRNA using Trizol. The goal of our analysis was not to quantify differences in expression levels per se, but rather to look for the complete absence of any gene expression. We deemed it equally informative and more cost-effective to increase the number of reads sequenced for each of the four life stage libraries, rather than produce independent biological replicates for each life-stage. Pooled samples, especially when the number of reads is high, have been shown to improve the power to detect gene expression of transcripts present at low abundances (Takele Assefa et al., 2020). Our use of pooled samples here, sequenced to produce a relatively high number of reads (>20 million each), should have facilitated an accurate representation of binary gene expression (expressed or not expressed), even for genes expressed at low levels.

After RNA extraction, we next prepared sequencing libraries following the Illumina TruSeq protocol, incorporating Covaris shearing as an alternative to chemical shearing and excluding the cDNA DSN normalization step. Paired-end sequencing was performed on a HiSeq2000 platform. We mapped the resulting reads from each sample to the *Cx. quinquefasciatus* genome with STAR v. 2.5.2 (Dobin et al., 2013, Dobin and Gingeras, 2015) as implemented in RSEM v. 1.3.1 (Li and Dewey, 2011). RSEM was then used to calculate the Transcripts per Kilobase Million (TPM) and Fragments Per Kilobase of transcript per Million mapped reads (FPKM) for all genes in each of the four samples. We considered any measure of either TPM or FPKM above zero as evidence of possible gene expression.

### Exogenous Influences on Circadian Rhythms

To complement our genetic analyses, we wanted to assess whether *Cx. molestus* displays behaviors that can be influenced by exogenous influences, specifically variation in light cues. To do this, we utilized a lab colony of *Cx. molestus* that originates from adult females collected in a New York City, NY residential basement in December 2010 (Price and Fonseca, 2015). It has been continuously maintained in various labs from that time to the present. Adults in our lab colony are kept in 60 cm^3^ screened flight cages (BugDorm-2120 Insect Rearing Tent), where they are allowed to mate freely. They are given access to an 8% sucrose solution *ad libitum*. As *Cx. molestus* are autogenous, females do not require a blood meal before they will lay eggs. To start a new generation, we collect egg rafts by placing a black takeout food tray (henceforth, “oviposition trays”) with 400 mL of dechlorinated water into the cages for 48 hours. We then remove these trays and allow the eggs to hatch. For colony maintenance, early instar (first and second) larvae are transferred from the oviposition trays to 600 mL of dechlorinated water in 1 L white plastic deli containers. We feed larvae a diet of TetraMin tropical fish flakes approximately once per week, with the amount varying depending on larvae size and density. Individuals are allowed to pupate in these containers and upon emergence, adults are transferred to the aforementioned flight cages. All life stages are maintained at ambient temperatures in a windowless room with no fixed light cycle.

Over three separate trials, we obtained 17 egg rafts from our maintenance colony as described above. After hatching, we placed five to fifteen larvae from each family into a 100 mL glass jar with 50 mL of dechlorinated water. These jars were then kept in a TriTech™ Research Digitherm^®^ 38-liter Heating/Cooling Incubator, with optional light blocking door coating and seal (https://www.tritechresearch.com/DT2-MP-47L.html) at one of four environmental conditions: 12:12 light:dark (lights on at 06:00, lights off at 18:00), 12:12 dark:light (lights off at 06:00, lights on at 18:00), 24 hours of light, or 24 hours of dark. All four incubators were kept at 23 °C and larvae were fed as described above with the amount varying based on larval size and density.

After pupation, we transferred individual pupae into polystyrene ‘wide’ drosophila vials (Genesee Scientific) that contained approximately 25 mL of dechlorinated water. These vials were then moved into a dark chamber with a light regime of 12:12 light:dark (lights on at 06:00, lights off at 18:00). During the dark period, a red light would come on once every hour for one minute in order to help detect any emergence activity. Adult emergence was filmed using a YI 1080p home security camera with night vision capability. Video technology such as this has been used to record mosquito behavior and activity in laboratory settings previously, including under light-dark and constant dark conditions (e.g. Araujo et al., 2020). We focused on the transition event (eclosion) between the pupal and adult states because it is relatively quick and discrete, and is easily noted in video footage (see Fig. S1).

We reviewed all the video recordings and documented the times of all emergences that occurred during the trials, as evidenced by observing the mosquito emerging from its pupal case or else the first point at which an adult was observed treading on the surface of the water (Fig. S1). For the purposes of analysis, we converted minutes into decimals (i.e. an adult emergence at 12:30 was recorded at 12.50). The Rayleigh Test was used to test for uniformity of the data (i.e. if adult emergence occurred at a certain time of the day). Our null hypothesis was that there was no pattern displayed in the adult eclosion of *Cx. molestus*. To test for differences between treatments, we used the Watson-Williams Test of Homogeneity of Means, performing all pairwise comparisons. All statistical analyses were done in R v. 4.0.2 (R Core Team, 2020), utilizing the ‘circular’ package (Agostinelli and Lund, 2017), and statistical significance was considered at α ≤ 0.05.

### Photoperiodically induced Dormancy (Quiescence or Diapause)

A true diapause state in female *Culex* mosquitoes is typically assayed by removing the ovaries and measuring the length of the follicles (e.g. Eldridge, 1968; Spielman and Wong, 1973; Sim and Denlinger, 2008; Meuti et al., 2015). All North American *Cx. molestus* populations examined in this manner have been shown to undergo ‘normal’ ovarian development when reared in short-day conditions (Spielman and Wong, 1973; Nelms et al., 2013). However, some members of the *Culex pipiens* species complex, such as *Cx. australicus*, are known to enter a quiescent state during inclement weather, that while not true diapause, nonetheless appears to be an adaptation to variable climates (Dobrotworsky and Drummond, 1953; Dobrotworsky, 1967). It is possible that a dormant state more akin to quiescence than to diapause (and independent of ovarian arrest) could be induced in *Cx. molestus* in response to photoperiodic cues (Diniz et al., 2017). Therefore, we wanted to assess whether our New York City *Cx. molestus* line expressed any degree of ecologically-relevant dormancy (e.g. ‘quiescence’) in response to a short photoperiod.

To do this, we collected egg rafts from 12 different females from our stock colony using oviposition trays in the manner described above. Each raft was isolated as a single family group and retained at 18 °C with a 12:12 Light:Dark (L:D; 7:00 - 19:00) photoperiod. Prior to their hatching, we gave each family group 100 μl of a solution made from vigorously blending 100 milligrams of fish flakes into 10 mL of dechlorinated water. 24 hours after hatching, we split each family group into approximately equivalent (numerically) groups that were then assigned to one of two photoperiod conditions, ‘long day’ and ‘short day’. Larvae placed in long-day conditions were kept in an 18:6 L:D photoperiod (4:00 - 22:00) at 18 °C, whereas larvae in short-day conditions were kept in a 6:18 L:D photoperiod (10:00-16:00), also at 18 °C. All egg, larval and pupal environmental conditions were maintained utilizing a TriTech™ Research Digitherm^®^ 38-liter Heating/Cooling Incubator, with optional light blocking door coating and seal (https://www.tritechresearch.com/DT2-MP-47L.html). First and second instar larvae were given 200 μl of fish flake feeding solution daily, third instar larvae were given 400 μl daily, and forth instar larvae were given 800 μl daily.

Upon pupation, individual mosquitoes were placed in a 100 mL glass jar with 50 mL of dechlorinated water. These pupae were maintained in the same environmental conditions as the larvae, depending on treatment group (long day vs. short day). When they eclosed (emerged as adults), we moved adult females to a flight cage with virgin males from a separate, dedicated mating pool. Males from this mating pool were maintained in the long day photoperiod at 20 °C from hatching to adulthood. The slightly higher temperature was to ensure that males reached sexual maturity before their interaction with experimental females. Males were at least three days old prior to the introduction of experimental females to ensure these males were sexually active (Vinogradova, 2000). Furthermore, we maintained a ratio of two virgin males to each experimental female in each flight cage.

To maximize the likelihood of insemination, we kept females in the flight cages for 72 hours. These flight cages were maintained at long-day photoperiod (18:6), but ambient temperature (~25 °C ± 2 °C during the duration of the study). After 72 hours, females were placed individually into 100 mL glass jars with 50 mL of dechlorinated water to lay eggs. These jars were kept in long day conditions and at ambient temperature.

We examined multiple traits associated with reproductive activity in females. First, we compared the percentage of females from each treatment that laid eggs within ten days after being placed in an oviposition jar. Females *Cx. molestus* typically lay eggs within four to five days after eclosion when maintained at 25 °C and six to nine days when maintained at 20 °C (Vinogradova, 2000). In contrast, after diapause termination, the sister taxon *Cx. pipiens* generally requires ten days before ovarian follicle development and the laying of eggs (Tate and Vincent, 1936). For individual females, we recorded the time to lay eggs (checked every 12 hours at 9:00 and 21:00), and the number of eggs laid. To count the number of eggs, we used a Nikon stereoscopic microscope (model C-PS) with a Gosky 10X microscope Smartphone Camera Adaptor to photograph each egg raft at 40x magnification. The eggs within each image were then marked and counted digitally using the program ImageJ v. 1.53a (Schneider et al., 2012). Because a female’s size can influence the number of eggs she lays (Vinogradova, 2000), we also measured wing length as a proxy for size using a metric miniscale (https://www.bioquip.com/search/DispProduct.asp?pid=4828E). Females were killed with ethyl acetate prior to wing measurement.

## RESULTS

### The gene, *Helicase domino*, harbors a Cx. molestus-specific structural variant

Our analysis of potential ‘large effect’ variants in *Culex* circadian genes identified 33 candidate genes (Table S4). However, upon further examination and confirmation, 32 of these genes were either not specific to the *Cx. molestus* samples examined, or else were mis-characterized due to apparent inaccuracies in the *Cx. quinquefasciatus* genome annotation. The single gene that was correctly annotated, present in only *Cx. molestus* samples, and absent from all *Cx. pipiens* samples, was a nine nucleotide, in-frame deletion in the fifth exon of the Helicase domino (*dom*) gene (Figure 1). This variant was present in *Cx. molestus* RNA-seq and genomic data from New York City, Germany, and Belarus, but absent in examined *Cx. pipiens* samples from similar, geographically proximate locations.

**Figure 1.**
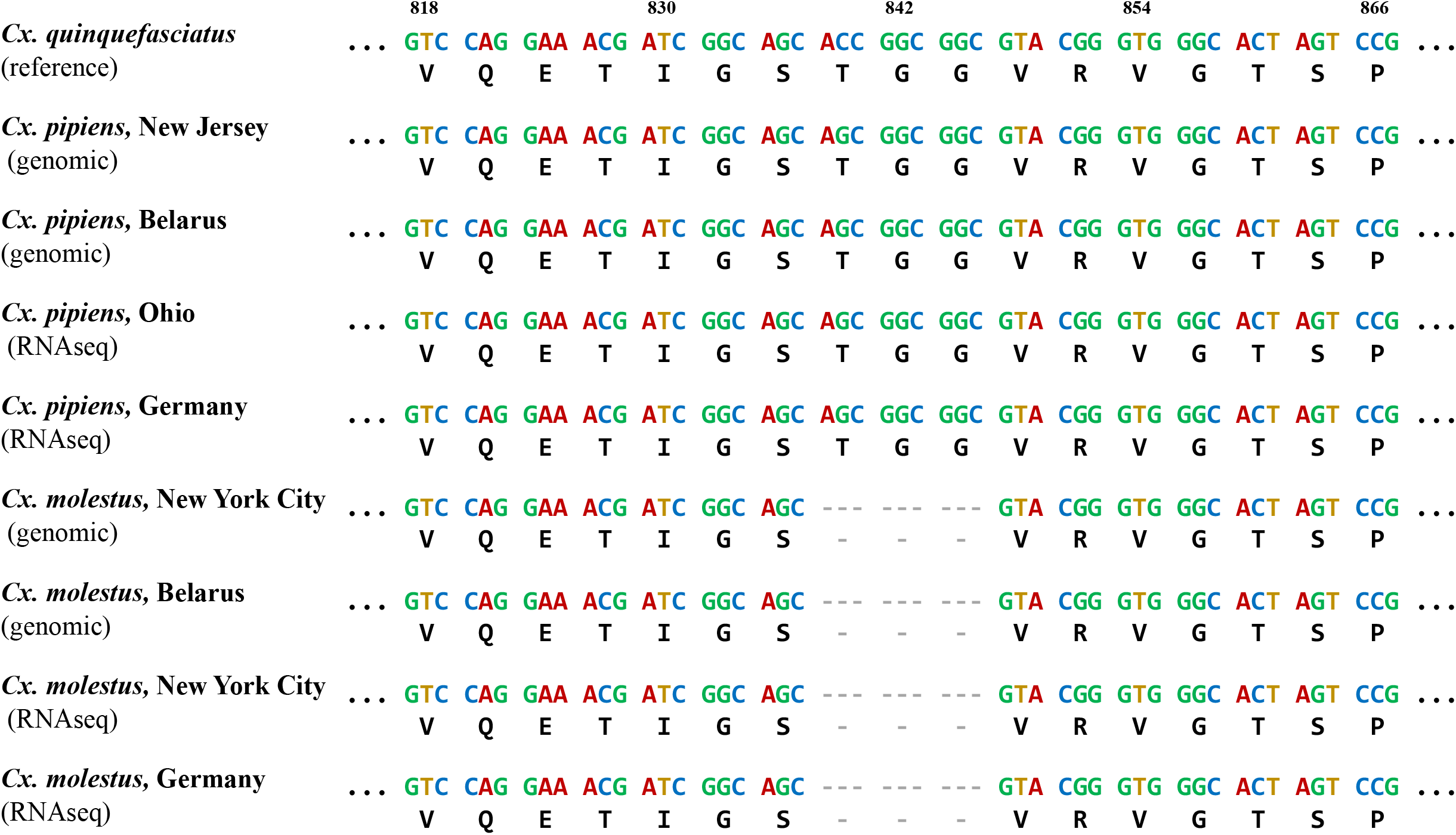
An alignment showing the nine nucleotide, in-frame deletion in the *Cx. molestus* Helicase domino (*dom*) gene. This deletion is located in exon 5, between nucleotides 838 to 846 (in relation to the annotated gene sequence in *Cx. quinquefasciatus*). The one letter codes for the corresponding amino acids are also given below each nucleotide sequence. The sample taxa, geographic origin, and data type are given to the left of each nucleotide/amino acid combination.

### Additional circadian genes harbor non-synonymous single nucleotide polymorphisms (SNPs)

Based on previous analysis of non-synonymous divergence between *Cx. molestus* and *Cx. pipiens* (Price and Fonseca, 2015), we examined 49 *Culex* circadian genes for derived amino acid changes relative to *Cx. quinquefasciatus*. Of these, eight genes had one derived amino acid change (missense mutation) in both the examined New York City and Belarussian *Cx. molestus* genome samples, and which were absent in the New Jersey and Belarussian *Cx. pipiens* genome samples (Tables 1, S5). As annotated in *Cx. quinquefasciatus*, these genes were: ‘calmodulin binding transcription activator 2’, ‘sodium chloride dependent amino acid transporter’, ‘dna photolyase’, ‘calmodulin-binding protein trpl’, ‘ultraviolet-sensitive opsin’, ‘glycogen synthase kinase 3’, ‘phospholipase c’, and a conserved hypothetical protein. There were 12 genes that had derived amino acid changes in both examined *Cx. pipiens* samples, with many genes harboring more than one missense mutation (21 total derived amino acid changes in *Cx. pipiens*).

**Table 1.** *Culex* circadian genes found to harbor a fixed, derived amino acid change (missense mutation) in both a New York City (NYC, USA) and Minsk (Belarus) *Cx. molestus* sample. The derived state was determined in relations to the annotated reference of *Cx. quinquefasciatus*. The *Cx. pipiens* samples derive from New Jersey (NJ, USA) and Minsk (Belarus). For more details, see Table S5.

| *Cx. quinque.*<br>Reference<br>Gene ID | Length<br>of<br>Gene<br>(AAs) | Amino<br>Acids<br>Analyzed | NYC<br>(*m.*) | Belarus<br>(*m.*) | Both<br>*molestus* | NJ<br>(*p.*) | Belarus<br>(*p.*) | Both<br>*pipiens* | *Cx. quinque.* Gene<br>Annotation |
| --- | --- | --- | --- | --- | --- | --- | --- | --- | --- |
| CPIJ003689 | 244 | 213 | 0 | 0 | 1 | 0 | 0 | 1 | calmodulin binding<br>transcription<br>activator 2 |
| CPIJ015063 | 618 | 605 | 1 | 1 | 1 | 0 | 0 | 1 | sodium chloride<br>dependent amino<br>acid transporter |
| CPIJ009455 | 499 | 417 | 1 | 1 | 1 | 2 | 1 | 0 | dna photolyase |
| CPIJ005741 | 1295 | 699 | 0 | 0 | 1 | 0 | 3 | 3 | calmodulin-binding protein trpl |
| CPIJ005000 | 378 | 261 | 1 | 1 | 1 | 0 | 0 | 0 | ultraviolet-sensitive opsin |
| CPIJ006114 | 503 | 495 | 1 | 0 | 1 | 2 | 0 | 0 | glycogen synthase<br>kinase 3 |
| CPIJ000778 | 1069 | 870 | 2 | 1 | 1 | 1 | 0 | 0 | phospholipase c |
| CPIJ016941 | 577 | 417 | 0 | 0 | 1 | 0 | 1 | 0 | conserved<br>hypothetical<br>protein |

### *Culex* circadian gene expression in *Cx. molestus*

Our sequencing of four pooled libraries each constituting a distinct *Cx. molestus* life-stage (larvae, pre-pupae, pupae, and adult) resulted in over 21 million read pairs per library (Larvae: 24.7M; Pre-pupae: 23.6M; Pupae: 21.1M; Adult: 22.4M). This sequencing data is deposited in the NCBI SRA database (accession numbers SRRXXXXXXX-SRRXXXXXXX). Of the 154 identified *Culex* circadian genes, all but four had evidence of expression in at least one *Cx. molestus* life history stage (Tables 2, S6). These four genes are annotated in the *Cx. quinquefasciatus* genome as ‘AMP dependent ligase’, ‘tubulin beta-3 chain’, ‘Dual specificity tyrosine-phosphorylation-regulated kinase’, and an uncharacterized protein. This last gene appears most similar to ‘E3 ubiquitin-protein ligase TRIP12’ in *Drosophila melanogaster*. Six other genes were very lowly expressed (TPM & FPKM < 1) in only one of the four examined life stages (Tables 2, S6).

**Table 2.** *Culex* circadian genes that were either not expressed in any *Cx. molestus* life history stage (Transcripts per Kilobase Million [TPM] = 0, bolded rows), or else in which TPM was < 1.0 for only one life stage. For more details, see Table S6.

| Gene ID | Larvae TPM | Pre-pupae TPM | Pupae TPM | Adult TPM | *Cx. quinque.* Gene Annotation |
| --- | --- | --- | --- | --- | --- |
| CPIJ002636 | 0 | 0 | 0.15 | 0 | protein kinase |
| CPIJ005003 | 0 | 0.17 | 0 | 0 | Predicted protein |
| **CPIJ010494** | **0** | **0** | **0** | **0** | **AMP dependent ligase** |
| CPIJ011756 | 0 | 0 | 0.57 | 0 | uncharacterized protein |
| **CPIJ012634** | **0** | **0** | **0** | **0** | **tubulin beta-3 chain** |
| **CPIJ014803** | **0** | **0** | **0** | **0** | **Dual specificity tyrosine-phosphorylation-regulated kinase** |
| **CPIJ014875** | **0** | **0** | **0** | **0** | **uncharacterized protein** |
| CPIJ015063 | 0 | 0 | 0 | 0.15 | sodium/chloride dependent amino acid transporter |
| CPIJ015933 | 0 | 0 | 0 | 0.22 | expressed protein |
| CPIJ017982 | 0 | 0 | 0 | 0.46 | casein kinase I isoform alpha |

### Exogenously-influenced circadian rhythms are retained

The Rayleigh test indicated that all treatments displayed clustering in the data (Figure 2; LD: 0.6042, P < 0.001; DL: 0.3413, P < 0.001, LL: 0.2051, 0.009, DD: 0.2301, P = 0.0045). The mean emergence time for *Cx. molestus* adult eclosion for individuals reared in 12:12 light:dark (‘LD’, lights on at 06:00, lights off at 18:00) was 21:58 (SD±1:00), approximately four hours after the onset of the dark cycle during larval development. For individuals reared in 12:12 dark:light (‘DL’) the mean was 10:08 (SD±1:28), again approximately four hours after the onset of the dark cycle during larval development. The mean adult eclosion time for individuals reared in constant light (‘LL’, 24 hours light) was 20:15 (SD±1:47), and for individuals reared in constant dark (‘DD’, 24 hours dark) was 17:50 (SD±1:43). In our pairwise comparisons, we observed a statistically significant difference between our LD and DL treatments (W_2_=75.199, p < 0.001), between our LD and LL treatments (W_2_=22.821, p < 0.001), between out LD and DD treatments (W_2_=32.759, p < 0.001), between our DL and LL treatments (W_2_=28.856, p < 0.001), and between our DL and DD treatments (W_2_=20.602, p < 0.001). Between our LL and DD treatments, there was no significant difference (W_2_=2.2454, p = 0.3254).

**Figure 2.**
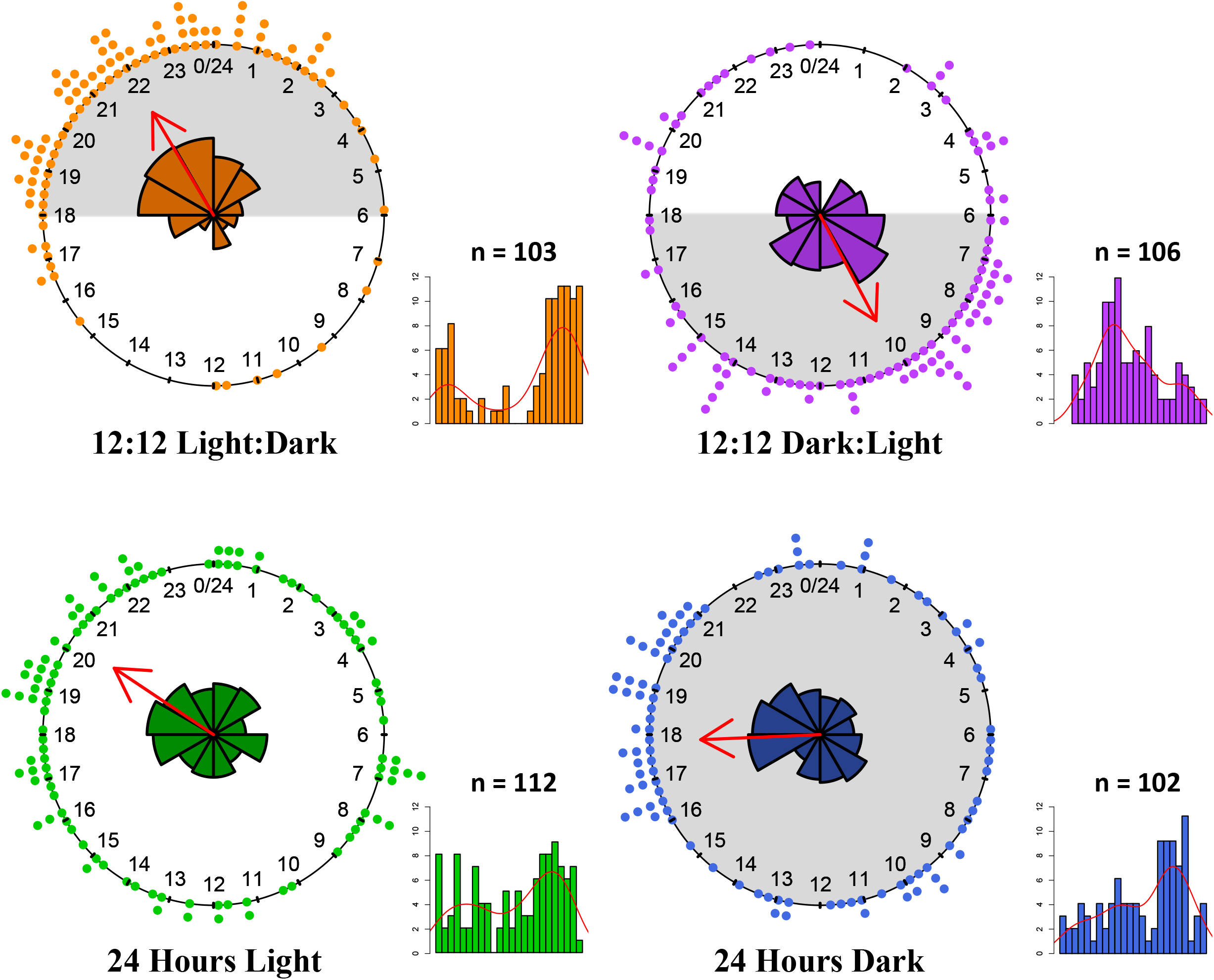
The distribution of adult eclosion times for each of our four treatments (12hours:12hours light:dark; 12 hours:12hours dark:light; 24 hours light; 24 hours dark). The dots around each compass represent one emergence on a 24-hour clock. The red arrows within each compass indicate the mean emergence time. In the middle of each compass is a rose diagram with 12 bins showing the circular distribution of emergence times. Grey areas of the compasses indicate periods of time when the larvae for each treatment were in darkness. To the lower right-hand side of each compass is a linear histogram of emergence times divided across 24 hours (from 0:00 [midnight] to 23:59). The red lines indicate the density distribution of emergences. The numbers above each liner histogram indicate sample sizes.

### Absence of photoperiodically induced dormancy

Of the 109 females set up for oviposition from those reared in long-day conditions, 102 laid eggs (93.6%), and of the 101 females set up for oviposition from those reared in short-day conditions 94 laid eggs (93.1%). On average, a female reared in long day conditions took 38.4 hours (SD±33.8) to lay eggs, and females reared in short day conditions took 47.9 hours (SD±45.1). The variance between the two groups was not statistically different (Levene’s Test, F_1,192_= 1.79, p= 0.1825), so we performed a t-test assuming equal variances. The results of this test indicated that there was no statistically significant difference in the time it took for females to lay eggs between treatments (t_192_= −1.6683, p = 0.097).

On average, females reared in long-day conditions laid 81.4 eggs (SD±18.3), whereas females reared in short-day conditions laid an average of 97.8 eggs (SD±15.8). There was a statistically significant difference between the two treatments (t_181.32_=-6.5394, p < 0.001). An ANOVA additionally showed that these differences between treatment were significant (F_1,192_= 42.21, p < 0.001). However, there is a strong intraspecific relationship between mosquito size and number of eggs laid (Vinogradova, 2000), and we observed that on average female mosquitoes reared in long day conditions had a wing length of 4.38mm (SD±0.21), whereas females reared in short day conditions had an average wing length of 4.68mm (SD±0.16). This difference was significant (t_174.67_ = −10.851, p < 0.001). When we controlled for these observed differences in size using an ANCOVA, there was no difference between the treatments (F_1,181_=0.350, p = 0.555). Figure 3 shows the relationship between wing size and the number of eggs laid for both treatment groups.

**Figure 3.**
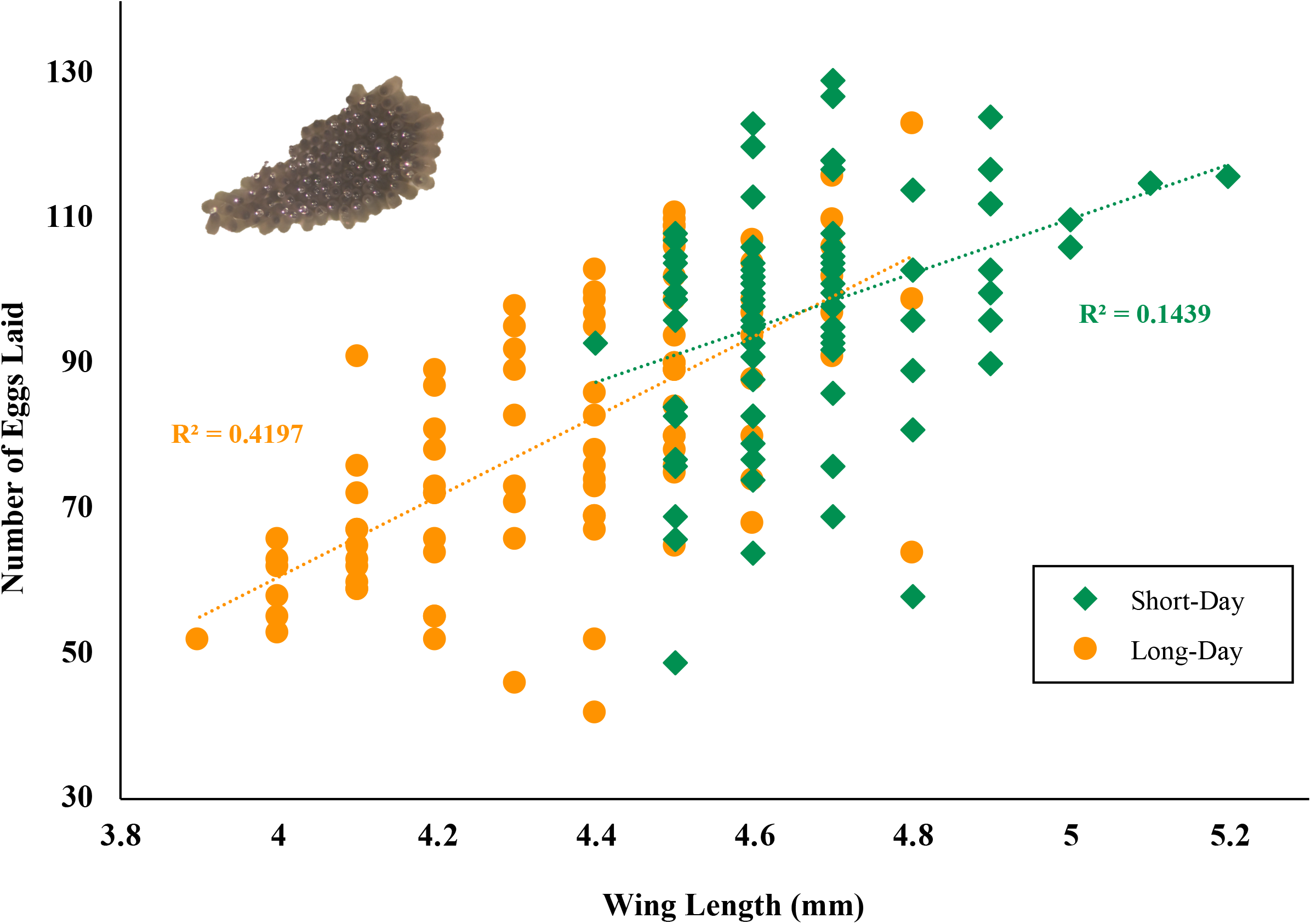
Scatter plot indicating the relationship between adult female wing size and the number of eggs laid. Data from females reared in short day conditions (6:18 Light:Dark) are indicated with green diamonds, and data from females reared in long day conditions (18:6 Light:Dark) are indicated with orange circles. The linear regression lines and R^2^ values for each dataset are also given. An example egg raft is shown in the upper left corner of the figure.

## Discussion

Despite their ecological differences in habitat preference and diapause induction ability, the amount of genetic divergence in genes potentially influencing circadian rhythms and/or overwintering behavior between the *Cx. molestus* and *Cx. pipiens* samples examined here was minimal, with most genes having no non-synonymous differences. This agrees with the close, and often challenging, taxonomic relationship of these two mosquitoes (e.g. Smith and Fonseca, 2004; Fonseca et al., 2004; Aardema et al., 2020). Of the 154 *Culex* circadian genes we examined, only one harbored a *Cx. molestus-specific* structural variant. This gene, Helicase domino (*dom*), was found to have a nine nucleotide, in-frame deletion in all *Cx. molestus* samples surveyed. Given that in *Drosophila melanogaster*, the *dom* protein appears to have many other important functions (Ruhf et al., 2001; Liu et al., 2019), it is not surprising that the observed structural variant would not radically alter the sequence of this gene. However, it is still possible that the observed variant does have an influence on the expression of circadian rhythms and, correspondingly, diapause induction. In *D. melanogaster*, distinct splice variants of the *dom* gene dramatically impact circadian behaviors (Liu et al., 2019). In particular they affect the expression of the negative circadian regulators period (*per*) and timeless (*tim*). Interference of both these regulators via RNAi caused *Cx. pipiens* females that were reared in short day, diapause-inducing conditions to direct develop (Meuti et al., 2015). The effect that the nine-nucleotide deletion we observed in *Cx. molestus* has on the function of the *dom* protein, particularly in relation to *per* and *tim*, will require further investigation.

In addition to this structural variant, we also found eight genes that each harbored one derived, non-synonymous amino acid change (missense mutation) in *Cx. molestus* samples, but not *Cx. pipiens* samples. One of these genes encodes for the protein phospholipase c. In *D. melanogaster*, this gene (*norpA*, FBgn0262738) is predominately expressed in the eyes, and mutations in this gene ultimately affect the visual input pathway and circadian entrainment (Collins et al., 2004). This gene may also regulate splicing of the *per* gene, which could impact both circadian rhythms and diapause. Another gene observed to have a non-synonymous change in *Cx. molestus* was the glycogen synthase kinase 3. In *D. melanogaster* the protein encoded by this gene *(sgg*, FBgn0003371) can influence phosphorylation of *tim* and likely also impacts *per* (Martinek et al., 2001). Intriguingly, expression of another glycogen synthase gene in *Cx. pipiens* was shown to impact the regulation of glycogen and lipid storages during diapause, and it was deemed essential for survival during winter dormancy (King et al., 2020).

Among the genes that were not expressed in any assessed *Cx. molestus* life-stage, ‘AMP dependent ligase’, is perhaps most interesting. This gene appears to be homologous with the *Drosophila* gene ‘Very long-chain-fatty-acid--CoA ligase bubblegum’ (*bgm*, FBgn0027348), which is predominantly a metabolic gene that influences fatty acid and lipid metabolism, but which also regulates the circadian sleep/wake cycle (Thimgan et al., 2014). In bumblebees, this gene appears to be upregulated prior to the onset of diapause (when metabolic reserves are being accumulated in the body), and downregulated during the actual diapause period (Amsalem et al., 2015). More broadly, it has been found that many metabolic genes are differentially expressed between diapausing and non-diapausing *Cx. pipiens* females (Kang et al., 2016), indicating the potential importance of such genes for successful diapause. While the observed amino acid change in this gene could potentially impact *Cx. molestus*’ ability to enter a dormancy state, it may also correlate with the absence of diapause in this mosquito and the lack of a need to accumulate large metabolic reserves.

Our experimental assessment of dormant tendencies showed that in conditions reported to induce diapause in the sister taxon, *Cx. pipiens*, there were no clear reductions in the tendency to lay, time to lay, nor number of eggs laid (when corrected for size). Indeed, females reared in short-day conditions (6:18 L:D) during the larval stage actually laid more eggs than females reared in long-day conditions (18:6 L:D). This is the opposite trend we would predict if females reared in short-day conditions were induced towards any degree of reduced reproductive output by their rearing conditions. The simple explanation for the greater average number of eggs laid by short-day females is that they were generally larger. There is a strong positive correlation between *Culex* female size and the number of eggs laid (Vinogradova, 2000). However, why the females in this experiment should on average be larger is less clear. Perhaps the most parsimonious explanation is that the incubators set for short-day conditions were slightly cooler than those set for long-day conditions (due to the differences in the amount of time the lights were off/on respectively). Although we used internal thermometers to ensure the temperature settings were maintained at ~18 °C, slight but consistent deviations from this temperature may have been sufficient to affect the size of these mosquitoes. Cooler temperatures during the larval period result in larger adult mosquitoes (Vinogradova, 2000).

The complete absence of any reduced reproductive output in our experiment strongly suggests that this line of *Cx. molestus* lacks any ability to exhibit a photoperiodic inducible dormant state. However, the results from our circadian rhythms study indicate that *Cx. molestus* circadian behavior can be exogenously entrained by light cues (specifically the timing of the photoperiod) during the larval stage. This refutes our hypothesis that genetic changes in one or more *Cx. molestus* clock gene could simultaneously account for its inability to enter diapause and produce a reduction or loss of entrainable circadian rhythms. Our results clearly show that this taxon maintains photoperiodic perception and some degree of circadian entrainment influenced by this perception. However, as we only investigated photoperiodic entrainment within a single *Cx. molestus* population, it remains unclear whether the degree and strength of this entrainment differs from other populations or taxa in the *Culex* genus.

One challenge of this study is the reliance on the available annotation of the closely related species *Cx. quinquefasciatus*. The original genome annotation predominately utilized automated gene annotation pipelines with some comparison to other Dipteran genomes available at the time. However, in our analysis of potential structural variants in *Culex* circadian genes, we observed multiple genes that appear incorrectly annotated in relation to the sequence of these genes assembled from RNA-seq data. The proportion of the discrepancies between the genome annotation and RNA-seq data that represent bioinformatic limitations versus those that are isoform differences or other biological variation remains to be determined. Regardless, this particular challenge likely limited an accurate characterization of structural variants present in the taxa examined here.

More fundamentally, this study only points towards potential genetic candidates for genetic changes within *Cx. molestus* that may correlate with its inability to enter dormant state in response to photoperiodic cues. We can provide no causative evidence that these changes contribute to this phenotype presently. Furthermore, the changes we have characterized are all likely to be derived within the *Culex pipiens* species complex. The justification for focusing on such derived genetic variation is that *Cx. molestus* is generally presumed to have evolved from a *Cx. pipiens* ancestor within relatively recent times (~10,000-80,000 years; Fonseca et al., 2004; Shaikevich, 2007). The evidence that *Cx. molestus* is the derived taxon is limited however, and it is interesting to note that our outgroup for comparisons, *Cx. quinquefasciatus*, also lacks an ability to enter a dormant state (Fonseca et al., 2004). If an absence of diapause is the ancestral condition in this mosquito group, and if *Cx. molestus* did not recently derive from the facultatively diapausing *Cx. pipiens*, then it is unlikely that derived genetic variation in contemporary populations is responsible for its inability to enter a dormant state in response to shortened photoperiods.

### Conclusions

We have shown that in the urban-adapted mosquito, *Culex molestus*, the ability to entrain circadian rhythms in response to exogenous light cues is decoupled from its inability to enter a photoperiodically induced dormancy state. Because it can mate in enclosed spaces and does not require a blood meal to reproduce, *Cx. molestus* is commonly maintained in entomology and disease vector laboratories worldwide. Given its wide usage in understanding mosquito biology, our results indicate that this taxon may be a valuable tool for exploring exogenously influenced phenotypes, particularly those that display a circadian rhythm. Greater knowledge of circadian rhythms in mosquitoes is critical for controlling and mitigating mosquito-vectored illnesses (Rund et al., 2016). Specifically, understanding how seasonal changes in photoperiod fine-tune daily mosquito behaviors could greatly improve our knowledge of transmission potential for specific pathogens and vectors. In this study, we uncovered a structural variant in the Helicase domino gene that segregates with the taxa examined here, and correspondingly with the ability to enter a diapause state. Given the substantial influences this gene appears to have on circadian rhythms in *Drosophila*, and given that circadian genes are known to greatly impact diapause induction in *Culex*, this gene represents a major target for follow-up studies. Additional genetic variation uncovered here specific to *Cx. molestus* also offers further opportunities to investigate the genetic underpinnings of the diapause trait. A better understanding of the genetic variation influencing diapause and circadian rhythms in *Culex pipiens* species complex mosquitoes more broadly may lead to improved vector control and a reduction in disease transmission.

## Acknowledgements

We would like to thank David Epstein for helping design and build the lighting mechanism. Prof. Bridgett vonHoldt graciously provided space for our diapause induction experiment. NRE was supported in part by a Bonnie Lustigman Research Fellowship. KS received funding for this project from the Wehner Student Research Program.

## Data availability

All raw RNA sequencing data have been deposited in NCBI’s GenBank under accession numbers SRRXXXXXXX-SRRXXXXXX.

## Supplementary information

**Table S1.** GO term IDs from *Drosophila melanogaster*, with corresponding definition (biological process). These terms were used to locate likely *Culex* circadian genes.

| Go Term | Definition |
| --- | --- |
| GO:0003052 | circadian regulation of systemic arterial blood pressure |
| GO:0003053 | circadian regulation of heart rate |
| GO:0007623 | circadian rhythm |
| GO:0009416 | response to light stimulus |
| GO:0009583 | detection of light stimulus |
| GO:0009648 | photoperiodism |
| GO:0009649 | entrainment of circadian clock |
| GO:0009793 | embryo development ending in seed dormancy |
| GO:0010162 | seed dormancy process |
| GO:0010231 | maintenance of seed dormancy |
| GO:0010840 | regulation of circadian sleep/wake cycle, wakefulness |
| GO:0022410 | circadian sleep/wake cycle process |
| GO:0022611 | dormancy process |
| GO:0042320 | regulation of circadian sleep/wake cycle, REM sleep |
| GO:0042321 | negative regulation of circadian sleep/wake cycle, sleep |
| GO:0042749 | regulation of circadian sleep/wake cycle |
| GO:0042752 | regulation of circadian rhythm |
| GO:0042753 | positive regulation of circadian rhythm |
| GO:0042754 | negative regulation of circadian rhythm |
| GO:0045187 | regulation of circadian sleep/wake cycle, sleep |
| GO:0045188 | regulation of circadian sleep/wake cycle, non-REM sleep |
| GO:0045938 | positive regulation of circadian sleep/wake cycle, sleep |
| GO:0048512 | circadian behavior |
| GO:0048571 | long-day photoperiodism |
| GO:0048572 | short-day photoperiodism |
| GO:0048573 | photoperiodism, flowering |
| GO:0048586 | regulation of long-day photoperiodism, flowering |
| GO:0048587 | regulation of short-day photoperiodism, flowering |
| GO:0050953 | sensory perception of light stimulus |
| GO:0050962 | detection of light stimulus involved in sensory perception |
| GO:0055115 | entry into diapause |
| GO:0061963 | regulation of entry into reproductive diapause |
| GO:0071482 | cellular response to light stimulus |
| GO:0071981 | exit from diapause |
| GO:0071982 | maintenance of diapause |
| GO:0097437 | maintenance of dormancy |
| GO:0097438 | exit from dormancy |
| GO:2000028 | regulation of photoperiodism, flowering |
| GO:2000033 | regulation of seed dormancy process |

**Table S2.** List of 154 *Culex* circadian genes (annotated in *Cx. quinquefasciatus*) investigated for structural variants, non-synonymous nucleotide changes (missense mutations) and absence of expression in *Cx. molestus*.

See spreadsheet

**Table S3.** List of *Culex* datasets used in this study with taxonomic designation, geographic origin of sample, data type, SRA number(s), and which analyses the data were used for. Also included is detailed information on the *Cx. quinquefasciatus* reference genome (with coding sequence annotation) used in this study.

See spreadsheet

**Table S4.** *Culex* circadian genes (annotated from *Cx. quinquefasciatus*) that contained potential ‘high-impact’ variants (see text for definition). Only one of these genes, Domino helicase, was determined to be a true biological variant, present in all examined *Cx. molestus* samples, and absent in all examined *Cx. pipiens* samples. The other 32 genes were either flagged due to absence in some *Cx. molestus* samples, presence in some/all *Cx. pipiens* samples, or inaccurate annotation of the *Cx. quinquefasciatus* genome. Also given are the gene ID, genome location of the gene, and the type of variant(s) observed.

See spreadsheet

**Table S5.** All *Culex* circadian genes (annotated from Cx. quinquefasciatus) that were previously found to have a Ka value greater than 0 in a comparison between *Cx. molestus* and *Cx. pipiens* (Price and Fonseca, 2015). The Amino Acid (AA) length of the annotated gene is given, as well as the number of amino acids analyzed here. Only amino acids that were present in all assessed genomes (i.e. no missing data or INDELs) and that which did not have a codon that spanned an intron were included. The number of derived amino acid changes observed in each sample or else multiple samples (depending on grouping of interest [taxon or geographic origin]). No observed amino acid is listed more than once. The derived amino acids of interest for this study are those that were observed in both North American and European *Cx. molestus* genomes, but neither geographically comparable *Cx. pipiens* genome (column F, highlighted in grey). Rows representing genes which contained an amino acid change in this category are bolded. For these genes we also include the likely *Drosophila melanogaster* homolog parent gene ID (based on our ‘blastp’ analysis), likely *D. melanogaster* homolog gene name, GO Terms associated with that gene, and these GO Term definitions.

See spreadsheet

**Table S6.** *Culex* circadian genes (annotated from *Cx. quinquefasciatus*) that were found to either exhibit no expression in any surveyed *Cx. molestus* life-stage (larvae, pre-pupae, pupae, or adult), or else in only one life-stage at a level below 1.0 Transcripts per Kilobase Million (TPM) and 1.0 Fragments Per Kilobase of transcript per Million mapped reads (FPKM). Also given are the *Cx. quinquefasciatus* gene annotations, likely *D. melanogaster* homolog parent gene ID (based on our ‘blastp’ analysis), likely *Drosophila* homolog gene name, GO Terms associated with that gene, and these GO Term definitions.

See spreadsheet

**Figure S1**. Focusing on the far left vile, this figure shows: A) *Cx. molestus* pupae before emergence B) *Cx. molestus* adult after emergence. The red arrows indicate the position of the pupae/adult respectively.

